# DENA: training an authentic neural network model using Nanopore sequencing data of Arabidopsis transcripts for detection and quantification of *N*^6^-methyladenosine on RNA

**DOI:** 10.1101/2021.12.29.474495

**Authors:** Hang Qin, Liang Ou, Jian Gao, Longxian Chen, Jiawei Wang, Pei Hao, Xuan Li

## Abstract

Models developed using Nanopore direct RNA sequencing data from *in vitro* synthetic RNA with all adenosine replaced by *N*^6^-methyladenosine (m^6^A), are likely distorted due to superimposed signals from saturated m^6^A residues. Here, we develop a neural network, *DENA*, for m^6^A quantification using the sequencing data of *in vivo* transcripts from Arabidopsis. DENA identifies 90% of miCLIP-detected m^6^A sites in Arabidopsis, and obtains modification rates in human consistent to those found by *SCARLET*, demonstrating its robustness across species. We sequence the transcriptome of two additional m^6^A-deficient Arabidopsis, *mtb* and *fip37-4*, using Nanopore and evaluate their single-nucleotide m^6^A profiles using *DENA*.

## Background

*N*^6^-methyladenosine (m^6^A) is the most abundant modification found in messenger RNA (mRNA) [1, 2]. Previous studies demonstrated that m^6^A affected RNA processes, such as pre-mRNA splicing[3, 4], RNA stability[5], and RNA localization[6]. Recently, Nanopore direct RNA sequencing (direct RNA-Seq) makes it possible to directly “visualize” signals of RNA m^6^A modification. Direct RNA-Seq studies on *in vitro* synthetic RNAs with adenosine replaced with *N*^*6*^-methyladenosine triphosphate substrates, observed signaling shifts between regular “A” residues and m^6^A residues[7] and increased base-calling errors around m^6^A residues[8]. Similarly, previous study also observed signaling shifts and base-calling errors between *in vivo* transcribed RNAs from wild-type and *ime4*Δ yeast mutant that lacked the m^6^A methyltransferase activity for RNA methylation[8]. Progress was made in locating m^6^A modification on RNAs using based-calling errors found in direct RNA-Seq reads from wild-type in comparison to those from m^6^A-deficient mutants. Taking advantage of this approach, several tools, such as *differr*[9], *DRUMMER*[10], *ELIGOS*[11], and *xPore*[12], were developed for m^6^A detection. However, this approach is hindered by the requirement of comparison of wild type and m^6^A-deficient mutants, and by the low sensitivity for hypomethylated m^6^A sites.

A more robust approach is to recognize m^6^A residues based on their unique electrical signal fingerprints in direct RNA-Seq data, which does not require m^6^A-deficient mutant and can pinpoint m^6^A residues on a given RNA molecule. For example, *MINES*[13] established a random forest model from ionic electrical value to call m^6^A events in sequence context of “AGACT”, “GGACA”, “GGACC”, and “GGACT”. Using direct RNA-Seq data generated from *in vitro* synthetic transcripts, several machine learning models were developed, such as *EpiNano* that uses support vector machines, and *Nanom6A* that uses Extreme Gradient Boosting[14]. Both can predict RNA m^6^A sites without the requirement of m^6^A-deficient mutants. However, some drawbacks in these methods hinder their performance on m^6^A prediction. For one, these methods were trained with the direct RNA-Seq data from *in vitro* synthetic transcripts, in which all “A” residues are replaced by m^6^A. Often the occurrences of clustered multiple m^6^A residues, especially consecutive m^6^A residues, may cause superimposed effect on signal readout. Thus, these models may not conform to those for studying m^6^A modification in *in vivo* transcribed RNA in live cells, and are limited in performance on natural samples. In addition, the training data generated from *in vitro* synthetic transcripts had limited numbers of variants in sequence context. For example, *Nanom6A* were trained with direct RNA-Seq data that contains 130 sites of “RRACH” motif that is substrate for *N*^6^-methyltransferase complex. They may not be sufficient to train the deep-learning models for m^6^A detection, as previous studies demonstrated that the performance on prediction can be improved on sophisticated deep neural networks[15, 16] using extended training data covering more diverse combinations of the five-residue motifs.

Here, we designed a novel neural network called *DENA* (**D**eeplearning **E**xplore **N**anopore m^6^**A**) by training on direct RNA-Seq data of *in vivo* transcribed mRNAs from wild-type and m^6^A-deficient *Arabidopsis thaliana* (*A.thaliana*). Compared to the *in vitro* synthetic RNA data, the *in vivo* transcribed RNA data are more productive in building reliable models on detecting m^6^A, as they: 1) do not contain clustered multiple m^6^A residues that may distort the m^6^A prediction model; and 2) extend coverage on more diverse combinations of the “RRACH” motifs in direct RNA-Seq data. *DENA* is shown to achieve accurate identification and quantification of m^6^A at single-nucleotide resolution in both *A.thaliana* and *Homo sapiens*. Importantly, *DENA* is able to detect m^6^A events on different isoforms of single genes (at single-isoform level), which is unavailable in previous methods. Furthermore, we evaluated *DENA* in two m^6^A-deficient *A.thaliana* mutants, *fip37-4* and *mtb*, and generated a rich resource for profiling m^6^A modification at single-nucleotide resolution for these *A.thaliana* lines. Our study established an approach to train with data containing naturally occurring m^6^A patterns from direct RNA-Seq sequencing of *in vivo* transcribed transcripts, and will provide a framework for identifying other types of RNA modifications using Nanopore direct RNA sequencing.

## Results and discussion

### Modeling with in *in vitro* transcripts of saturated m^6^A does not conform to that of m^6^A modifications in *in vivo* transcripts

Several models[8, 14] have been developed for RNA m^6^A identification utilizing the training data of *in vitro* transcripts. Nevertheless, there is a key limitation to consider. In the twelve possible five-mers of “RRACH” motif, ten of them contain at least two adenosine residues. However, all “A” residues are replaced with m^6^A residues in *in vitro* synthetic training data, causing the aggregation of multiple m^6^A residues, which is likely not to occur in *in vivo* transcripts from live cells. Thus, we investigate the difference between the cluster m^6^A residues in *in vitro* transcripts and the naturally occurring m^6^A residues in *in vivo* transcripts on electrical signals and base call accuracy around modification sites.

In *in vitro* synthetic transcripts, the shift of electrical signals[7] and the enrichment of the mismatches[8] around m^6^A residues have been observed in modified direct RNA-Seq reads relative to unmodified reads. We aligned the *in vitro* synthetic reads and *in vivo* transcribed reads to the corresponding reference, respectively (Figure 1a, d). We found the existence of clustered multiple mismatches within or around “RRACH” five-mers in m^6^A-modified reads of synthetic data (Figure 1a). For example, in two “AAACC” five-mers from *in vitro* synthetic sequences, we observed distinguishing signals and consecutive mismatches within the “AAACC” pattern and its surrounding “A” residues (Figure 1b). However, this situation was infrequent in direct RNA-Seq data of *in vivo* transcribed RNAs from *A.thaliana* (Figure 1d), and only sporadic distinction of signals and mismatches were observed around the m^6^A sites. For example, only single position occurs significant mismatches around two m^6^A sites, which are identified by *differr*[9] and contained in the m^6^A peak from MeRIP-Seq[17, 18] (Figure 1c, e). That is to say, the saturated m^6^A modification in *in vitro* synthetic transcripts led to different patterns of electrical signals and mismatches, relative to the m^6^A modification on *in vivo* transcripts. In addition, previous studies demonstrated that sophisticated neural networks can improve the performance on the prediction of chemical modifications[15, 16]. As mentioned in *introduction*, the direct RNA-Seq data from *in vitro* synthetic transcripts had a limited number of variants in sequence context, maybe insufficient for training deep-learning models for m^6^A detection.

**Figure 1.**
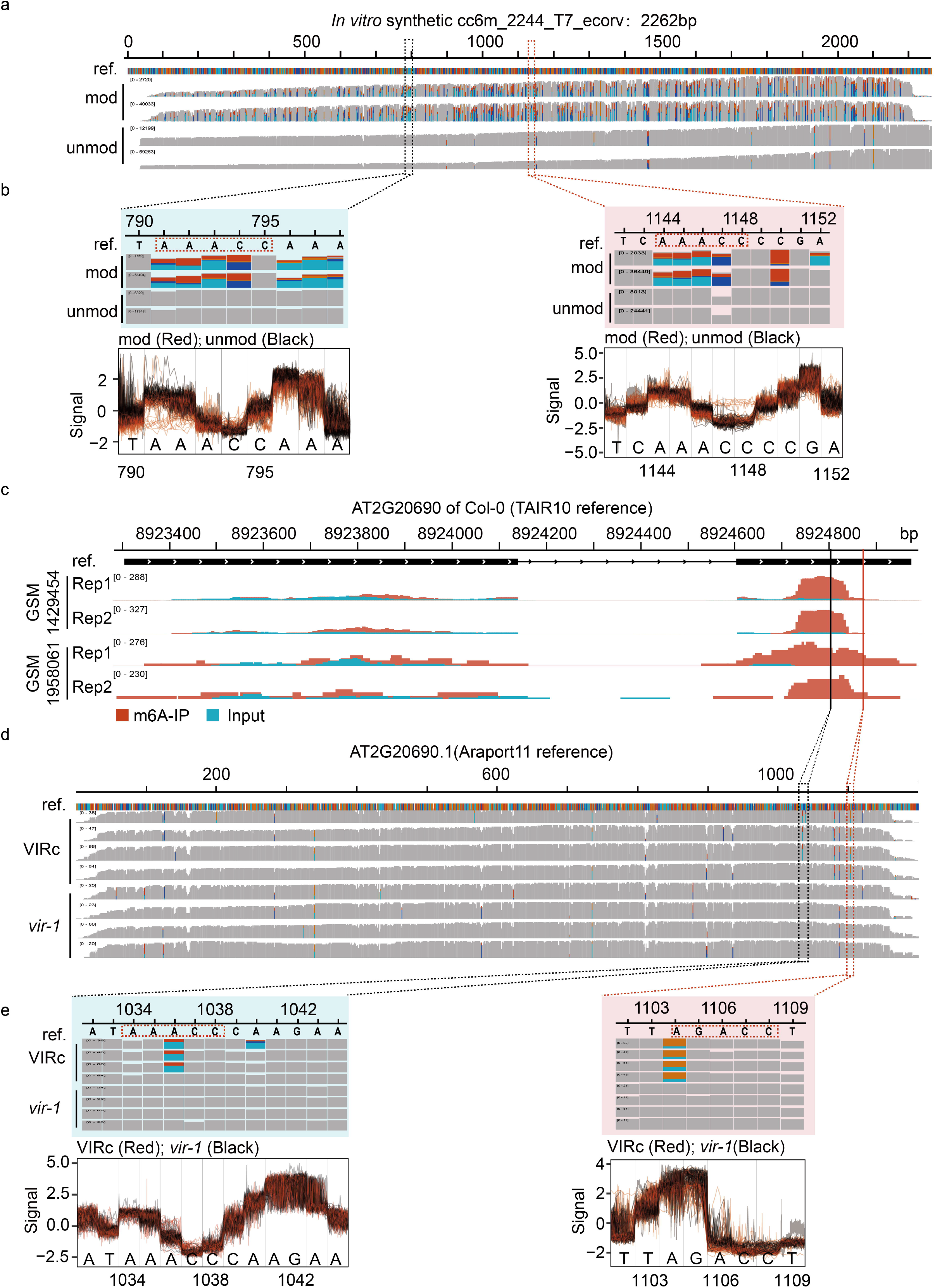
The distinction of electrical signals and mismatches between *in vitro* and *in vivo* transcripts. (a) The alignments are performed between the *in vitro* synthetic m^6^A-modified and unmodified reads and transcriptome reference. (b) The distributions of electrical signals and base-calling “errors” in *in vitro* reads at two “AAACC” sites. (c) The m^6^A peaks on AT2G20690 identified by two MeRIP-Seq datasets. (d) The alignments are generated by mapping the reads of VIRc and *vir-1* to Araport11 reference in the region of AT2G20690.1. (e) The distributions of electrical signals and base-calling “errors” at two m^6^A sites identified by *differr* tool in native direct RNA-Seq reads of VIRc and *vir-1*.

### Identification of m^6^A sites in *A.thaliana* for subsequent neural network training

In *A.thaliana*, m^6^A modification formed with recruitment of the m^6^A “writer” complex containing MTA (orthologue of human methyltransferase-like3, METTL3)[19], FIP37 (orthologue of human Wilms tumor 1-associated protein, WTAP)[17, 20], MTB (orthologue of human methyltransferase-like14, METTL14)[20], and VIRILIZER[20] *et al*.. We confirmed the significant decrease of m^6^A level in total RNA from two m^6^A-deficient mutants, *mtb* and *fip37-4*, using LC-MS/MS (*MATERIALS AND METHODS*), and performed direct RNA-Seq on the mRNAs extracted from Col-0, *mtb*, and *fip37-4*, respectively (Figure 2a). For each sample, over 1.8 million base-calling reads were aligned with the genome reference (TAIR10) of *A.thaliana* successfully (Additional file 1: Table S1), and their read-length were distributed within 800-1000 nucleotides (Additional file 1: Fig. S1a). In addition, the T-DNA insertions within FIP37 (AT3G54170) and MTB (AT4G09980) were also confirmed by direct RNA-Seq reads, respectively (Additional file 1: Fig. S1b).

**Figure 2.**
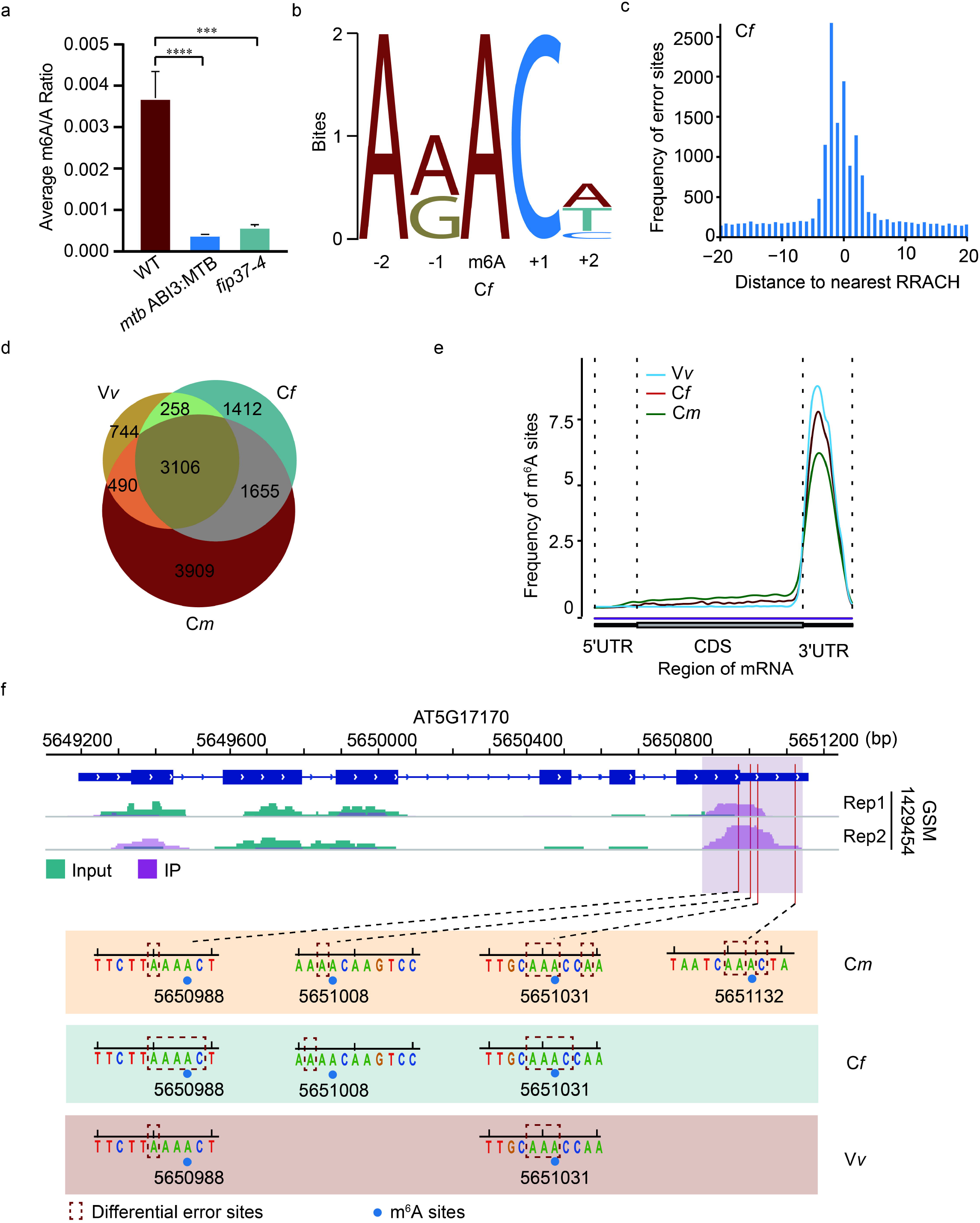
Quantification of m^6^A levels using LC-MS/MS and identification of m^6^A sites using *differr* from direct RNA-Seq reads. (a) LC-MS/MS quantify the ratio of “m^6^A” and “A” residues from total RNA of Col-0, *fip37-4*, and *mtb*, respectively. (b) The motif is enriched in C*f* group. (c) The distribution of distances between differential sites and its nearest “RRACH” motif in C*f* group. (d) Veen diagram shows the common m^6^A sites identified among C*m*, C*f* and V*v* groups. (e) The m^6^A distribution of three groups on transcripts, are strong enriched around the stop codon. (f) The m^6^A peaks detected by MeRIP-seq and the m^6^A sites identified by *differr* in ENH1 (AT5G17170) gene from three groups, respectively.

To obtain reliable m^6^A sites for subsequent neural network training, we used *differr* tool to compare the “mismatch” events in Col-0 with those in *mtb* (designated C*m*) and with those in *fip37* (designated C*f*) mutants, respectively. Searching for the motifs surrounding the differential sites detected by *differr*, we found the most frequently identified motifs in both C*f* and C*m* sets closely resembled the “RRACH” (R=A/G, H=A/C/U) motif established by Me-RIP-Seq in *A.thaliana*[17, 18, 21] (Figure 2b; Additional file 1: Fig. S1c). These differential sites mainly fell within five nucleotides of the nearest “RRACH” motif closest to them (Figure 2c; Additional file 1: Fig. S1d). In all, 6431 and 9160 m^6^A sites were detected in C*f* and C*m* using *differr*, respectively.

We further analyzed the direct RNA-Seq data from VIR-complemented (VIRc) and *vir-1* mutant (*vir-1*)[9] (designated V*v*), using the same procedure (Additional file 1: Fig. S1e, f). A total of 3106 m^6^A sites were common among C*m*, C*f*, and V*v* sets, accounting about 67.6% of m^6^A sites in the V*v* group (Figure 2d). A strong enrichment of m^6^A sites around the stop codon was observed in all three groups (Figure 2e). For example, *differr* detected four m^6^A sites in ENH1 (AT5G17170) and all of them fell within the established peaks from MeRIP-seq[18] (Figure 2f). Altogether, these results indicated these m^6^A sites identified by *differr* were reliable. The 3106 overlapped m^6^A sites were used for subsequent neural network training.

### Training a novel model with neural network for RNA m^6^A detection

The “error” events can occur at the adjacent bases of m^6^A sites due to the characteristic current blockade caused by m^6^A residues on RNA when performed direct RNA-Seq. We asked whether we were able to separate the *in vivo* direct RNA-Seq reads of *A.thaliana* basing on the error events that occurred within the nucleotide of m^6^A site and five nucleotides of upstream and downstream (Figure 3a). We extracted native m^6^A-modified and un-modified data of the 3106 m^6^A sites (shared by C*f*, C*m*, and V*v*) from VIRc and *vir-1*[9], respectively (Figure 1d; 3a). In all, we obtained the 2251835 matrices of features (mean, median, standard deviation, dwell time, and base quality) from the events of direct RNA-Seq reads, which contains 507066 in m^6^A-set and 1744769 in A-set (*MATERIALS AND METHODS*). We divided them into training and testing datasets at a ratio of 7:3 for training a neural network model, named *DENA* (Figure 3a). We evaluated the performance of our model, and for 12 possible five-mers of “RRACH” motif, the area under the curve (AUC) ranges from 0.90 to 0.97, and the accuracy from 0.83 to 0.93 in the testing dataset, indicating consistent performance among the twelve motifs (Figure 3b; Additional file 1: Fig. S2a; Additional file 1: Table S2). We applied *DENA* to the unmodified direct RNA-Seq data of the *in vitro* synthetic transcripts to evaluate its false positive. The background noise was found to be predominantly below 0.1, and almost all below 0.2 (Additional file 1: Fig. S2b). Thus, we designated the m^6^A sites as those above the baseline value 0.1, and the high-confidence m^6^A sites as those above 0.2 (s*ee METHODS section*).

**Figure 3.**
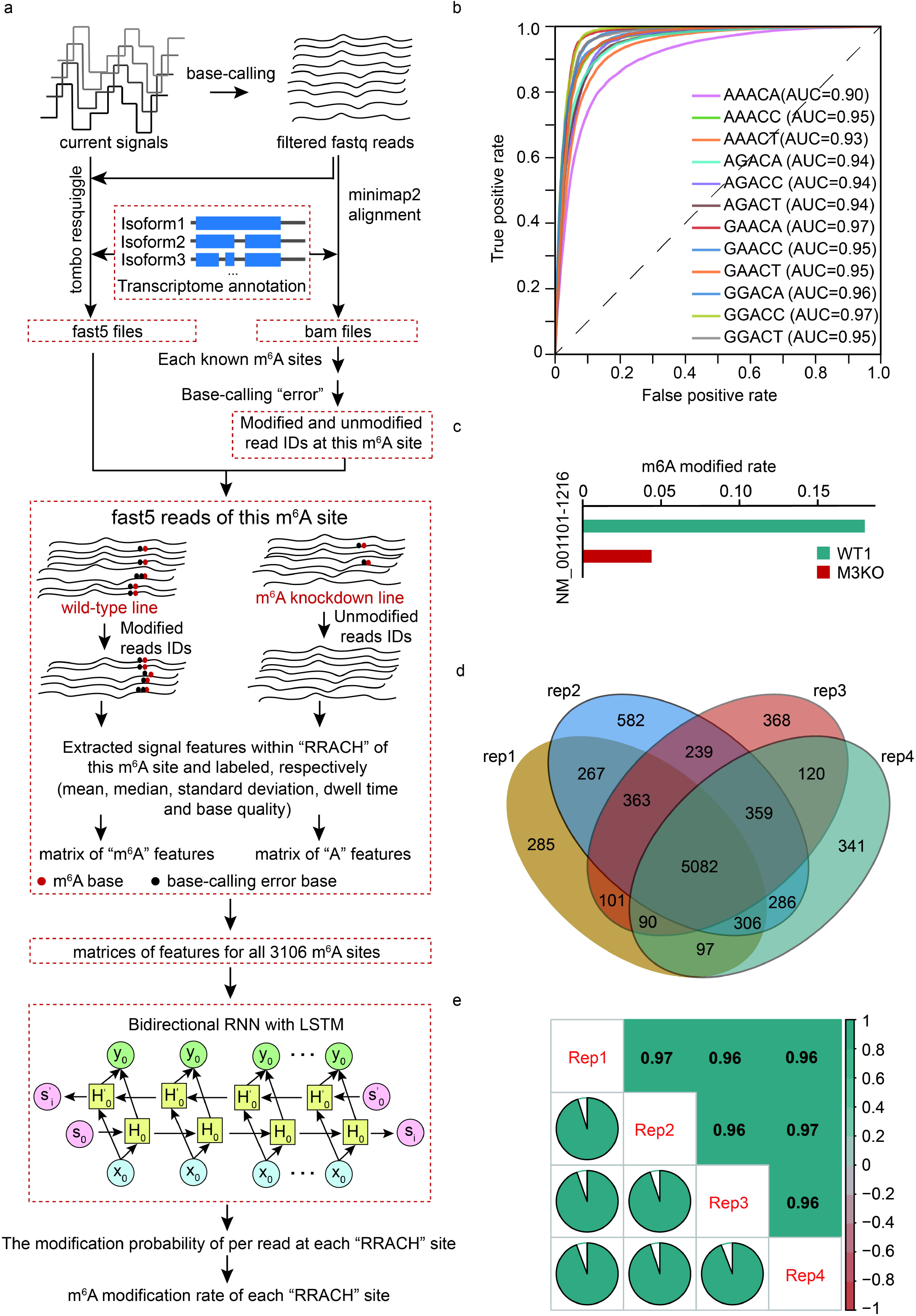
Extracted native training data and trained a neural network model. (a) Flowcharts illustrating the classification of modified and unmodified native reads and the procedures of training *DENA*. (b) The ROC curves of *DENA* in the 12 consensus sequences from “RRACH” motif. (c) Bar plots demonstrate the reduction of m^6^A rates identified by *DENA* at the ACTB-1217 site from direct RNA-Seq data of wild-type and METTL3-knockout human cells, respectively. (d) Venn diagram of the number of m^6^A sites identified by *DENA* from these sites that are supported at least 50 reads in all replicates of wild-type *A.thaliana*, respectively. (e) The cross-correlation coefficient of m^6^A rates identified by *DENA* among four replicates of wild-type *A.thaliana*.

To evaluate the ability of DENA in m^6^A detection and quantification, we performed predictions at the ACTB-1217 (NM_001101-1216) site, which was a known m^6^A site identified by the *SCARLET*[22] method previously, in direct RNA-Seq data of wild-type (WT) and METTL3 knockout (M3KO) human cells[23]. The modification rate at this site was 0.180 and 0.044 in WT and M3KO, respectively, suggesting a significant reduction of m^6^A modification in METTL3 knockout (Figure 3c). The *DENA*-identified rate in WT (0.180) was close to 0.21 that was identified by *SCARLET*, confirming the reliability of *DENA*.

To check whether *DENA* can reproduce m^6^A sites among replicated experiments, we tested DENA on the four replicates of wild-type *A.thaliana*[9]. To reduce the impact of the sequencing depth, we considered 22729 “RRACH” sites that were supported by at least 50 direct RNA-Seq reads in each of four replicates. A total of 6591, 7484, 6722 and 6681 m^6^A sites were obtained in the four replicates, respectively, and 5082 sites were common among them (Figure 3d). We computed the cross-correlation coefficient of the modification rate at these sites across four replicates, and obtained a correlation with 0.96 (Figure 3e). Altogether, these results confirmed the reliability of *DENA* for m^6^A detection and quantification.

### Validating the robustness of *DENA* in m^6^A detection in *A.thaliana* and human

To validate the ability of *DENA* in mapping m^6^A sites to transcriptome with single-nucleotide precision, we applied it to three published *A.thaliana* samples[9], including Col-0 (wild-type), VIRc, and *vir-1*, respectively. We scanned all “RRACH” sites on the transcriptome in three samples. 149136, 173730, and 166300 positions were supported by at least 50 direct RNA-Seq reads in Col-0, VIRc, and *vir-1*, respectively. A total of 46500 (31.18%), 48602 (27.98%), and 39241 (23.60%) m^6^A sites were identified by *DENA* in Col-0, VIRc, and *vir-1*, respectively, and 25,283 sites were overlapped across all samples (Figure 4a). A significant reduction of m^6^A modification rate was observed in *vir-1* compared to those in both Col-0 and VIRc (Figure 4b; Additional file 1: Fig. S3a). Importantly, the high correlation of modification rate was found between VIRc and Col-0, confirming the reliability of the m^6^A quantification using *DENA* (Figure 4c). The m^6^A sites identified by *DENA* enriched near the stop codon and 3′UTR in all samples (Additional file 1: Fig. S3b), agreeing with the distribution of m^6^A modification on mRNAs.

**Figure 4.**
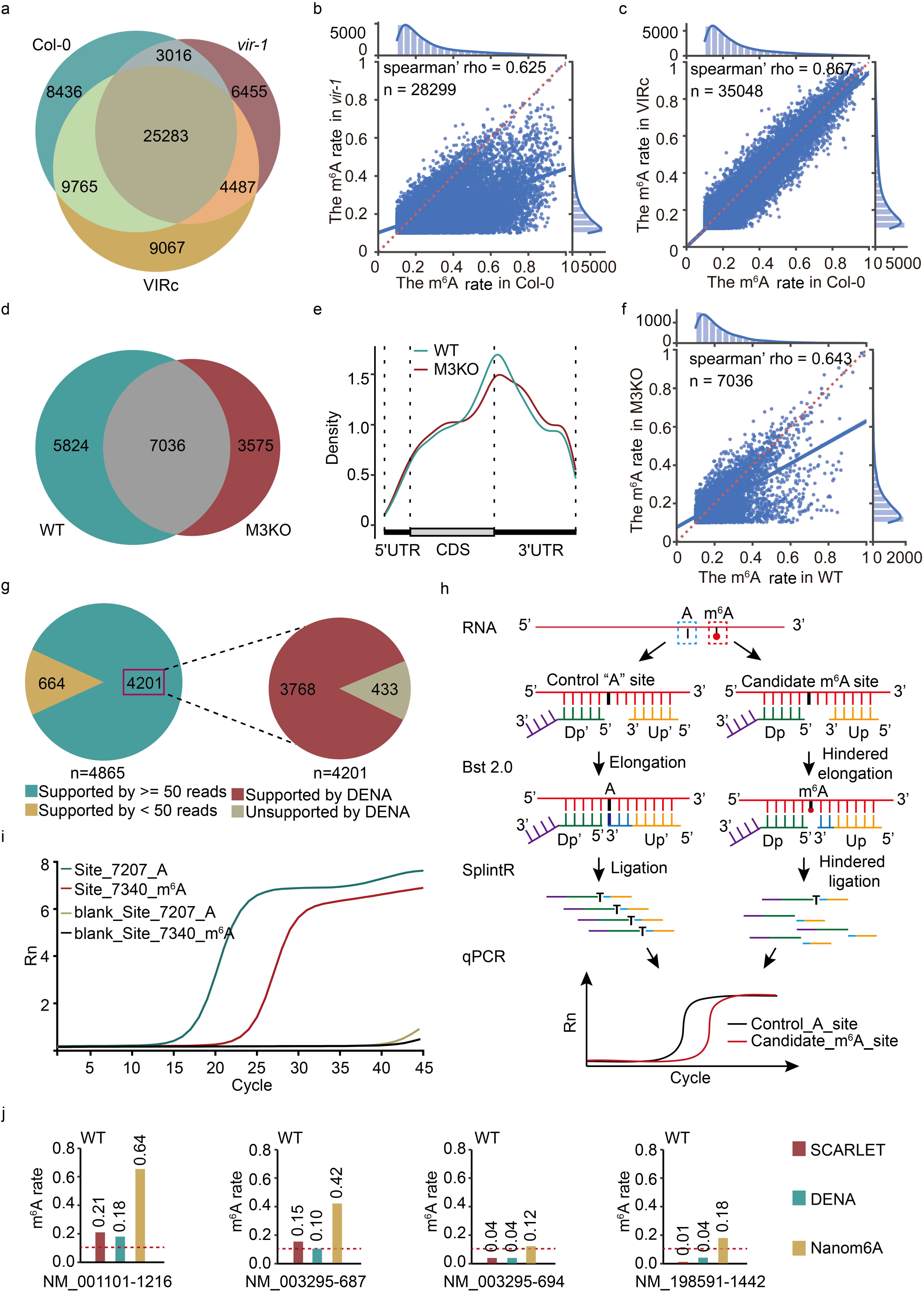
The m^6^A identification for *A.thaliana* and human using *DENA*. (a) Veen diagram shows the common m^6^A sites identified by *DENA* among Col-0, VIRc and *vir-1* samples. (b) Jointplot shows the correlation of m^6^A rates at 28299 overlapped sites between Col-0 and *vir-1*. (c) Jointplot shows the correlation of m^6^A modification rate at 35048 overlapped sites between Col-0 and VIRc. (d) Veen diagram shows the common m^6^A sites identified by *DENA* from wild-type (WT) and METTL3-knockout (M3KO) human cells. (e) The m^6^A distribution on transcripts in WT and M3KO, respectively. (f) Jointplot shows the correlation of m^6^A rates at 7036 overlapped sites between WT and M3KO. (g) Pie charts display the performance of *DENA* on 4201 miCLIP-identified m^6^A sites that are supported at least 50 reads, and 3768 sites out of them are regarded as m^6^A modification sites by *DENA*. (h) Flowcharts illustrating the identification of m6A sites using *Bst* 2.0 DNA polymerase Splint ligase with qPCR. Dp, and Up are down probe and up probe, respectively. (i) The real-time fluorescence curves produced at Site_7340 and Site_7207 on the mRNA of PRP8A, respectively. A is Site_7207 and m6A is Site_7340 on PRP8A. (j) Bar plot demonstrates consistency of m^6^A rate at the four known modified sites between *DENA* detection and *SCARLET* identification, and the significant overestimation in the prediction from *Nanom6A*.

We further assessed the *DENA* robustness on other species by applying it to human direct RNA-Seq data[23]. We obtained 12860 and 10611 m^6^A sites in WT and METTL3 knockout (M3KO) samples, respectively, and 7036 sites were common between them (Figure 4d). These m^6^A sites concentrated around the stop codon and 3′UTRs (Figure 4e). Notably, the modification rate in M3KO were significantly reduced relative to that in WT (Figure 4f).

We next compared *DENA* with *Nanom6A*[14] that was recently developed using the *in vitro* synthetic training data, using the 4685 miCLIP sites identified in previous work[9]. Among the 4685 m^6^A sites, 4201 had a sequencing depth of at least 50, of which about 90% (3768/4201) were detected as m^6^A modification by *DENA* (Figure 4g), a significant improvement compared with the 66% reported by *Nanom6A*. Meanwhile, we further randomly selected five *DENA*-predicted m^6^A sites (PRP8A_Site_7340, CURT1B_Site_1121, EMB1467_Site_2629, NACA3_Site_802 and RPL17B_Site_688) on mRNAs of Arabidopsis to identified by the *SELECT* experimental method[24], which utilizes the fact that m^6^A hinders both the single-base elongation activity of *Bst* DNA polymerase and the nick ligation efficiency of *SplintR* ligase (Figure 4h). For example, *DENA* identified a m^6^A site at 7340^th^ (Site_7340) nucleotide on the PRP8A transcript (AT1G80070.1). Using the nearby unmodified “A” (Site_7207) as a control to keep the same amount of RNA template, we performed an identification on this candidate m^6^A-modified site (Site_7340) with *SELECT* method. After performing simple qPCR-based quantification of the ligation products at Site_7340 site versus nearby Site_7207 site, we found that the Ct (threshold cycle) value at Site_7340 site was significantly increased relative to that at Site_7207, indicating that the amount of final ligation products formed at Site_7340 site was dramatically reduced compared to products formed at Site_7207, confirming the m^6^A modification at Site_7340 (Figure 4i). For the other four selected sites, we also detected the significant increase of Ct value at candidate m^6^A-modified site compared to that at nearby control “A” site, confirming the presence of m^6^A modification at these sites (Additional file 1: Fig. S3c). Therefore, these results confirmed the reliable prediction of m^6^A modification using *DENA* in Arabidopsis.

For human data, the m^6^A modification rate of several sites on two long non-coding RNAs (lncRNAs) (MALAT1 and TUG1) and three mRNAs (ACTB, TPT1 and BSG) were quantified by *SCARLET* method[22]. However, we found the Nanopore sequencing coverage of two lncRNAs were very low, both are fewer than 5 in all the datasets, which made it impossible to conduct a reliable analysis. We speculate about its cause and believe it is likely lncRNAs were not retained during the purification of mRNA using poly(T) before nanopore direct RNA-Sequencing, as the poly(A) structure may not be there for lncRNAs[25]. We thus compared *DENA* with *Nanom6A* at the known sites identified by *SCARLET* method on TPT1 (NM_003295), ACTB (NM_001101), and BSG (NM_198591) mRNAs[22]. The m^6^A rates quantified by *DENA* were more consistent with those by *SCARLET*, compared with those estimated by *Nanom6A* (Additional file 1: Table S3). For example, the NM_001101-1216 and NM_003295-687 sites were detected as m^6^A sites by *DENA* with 0.18 and 0.10 modification rates, respectively, which were close to the 0.21 and 0.15 quantified by *SCARLET*. However, m^6^A rates of 0.64 and 0.42 were obtained by *Nanom6A* at NM_001101-1216 and NM_003295-687, respectively (Figure 4j). For NM_003295-694 and NM_198591-1442 sites, *DNEA* obtained 0.04 and 0.04 modification rates, respectively, which were consistent with the value of 0.04 and 0.01 by *SCARLET*. However, *Nanom6A* returned 0.12 and 0.18 modification rates at NM_003295-694 and NM_198591-1442, respectively. These results suggested the significant overestimation of modification rate by *Nanom6A* relative to those by *SCARLET* (Figure 4j). We also investigated the m^6^A rate at these sites in M3KO cells using *DENA*. Notably, we observed significant reduction of modification rates at NM_001101-1216 and NM_003295-687 sites and slight decrease at NM_003295-694 and NM_198591-1442 compared to those in wild-type (Additional file 1: Fig. S3d). In addition, we further compared the predicted modification rates from *DENA* with those from other methods containing *xPore, LEAD-m*^*6*^*A-seq* and *Deoxyribozyme-based Method*, at the identified m^6^A sites in previous work[12, 26, 27] (Additional file 1: Table S3). Importantly, we can observe that the modification rates predicted by *DENA* are consistent with those identified by experimental methods at most sites. Concretely, the modification rates predicted by DENA at seven sites, ACTB_5527743, BSG_583239, BSG_583346, TPT1_45337310, TPT1_45337303, YTHDF2_28743593 and PARP1_226361173, are in general agreement with those verified by at least one experimental method, *SCARLET* and/or *LEAD-m*^*6*^*A-seq*. For example, NM_001618.4-3496, a negative site on PARP1 gene used to confirm the ability of *LEAD-m*^*6*^*A-seq* in previous work[26], was identified as non-m^6^A site with only 2% modification rate by *LEAD-m*^*6*^*A-seq*, which was comparable to the 7% modification rate predicted by *DENA*, while 44% modification rate was predicted by *Nanom6A*. In addition, the prediction of reduction in m^6^A modification rates in METTL3 knock-down mutant line using the three models, *Nanom6A, xPore* and *DENA*, were in general agreement (Additional file 1: Table S3).

Collectively, these results demonstrated the improvement and robustness of *DENA* in the detection and quantification of m^6^A modification in different organisms on transcriptome-wide.

### Identification of m^6^A sites on isoforms of single genes using *DENA*

Alternative splicing produces isoforms that include or exclude particular exons from the transcripts[3], representing an important mechanism for regulation of gene functions. Previous tools identified and quantified m^6^A modification by assigning them to genome reference[8, 11, 14]. Thus, they did not distinguish m^6^A modifications on isoforms produced by alternative RNA splicing. *DENA* was designed to detect and quantify m^6^A modification on isoforms from single genes (Figure 4a). For example, FNR2 (AT1G20020) encodes a leaf-type oxidoreductase in *A.thaliana* and is present in the chloroplast, having three isoforms based on Araport11 reference. We identified two, three, and two high-confidence m^6^A sites on the AT1G20020.1, AT1G20020.2, and AT1G20020.3 isoforms, respectively. This result indicated the differential m^6^A profiles in different isoforms of a single gene and affirmed the capability of *DENA* in the identification of m^6^A sites from different isoforms (Figure 5a).

**Figure 5.**
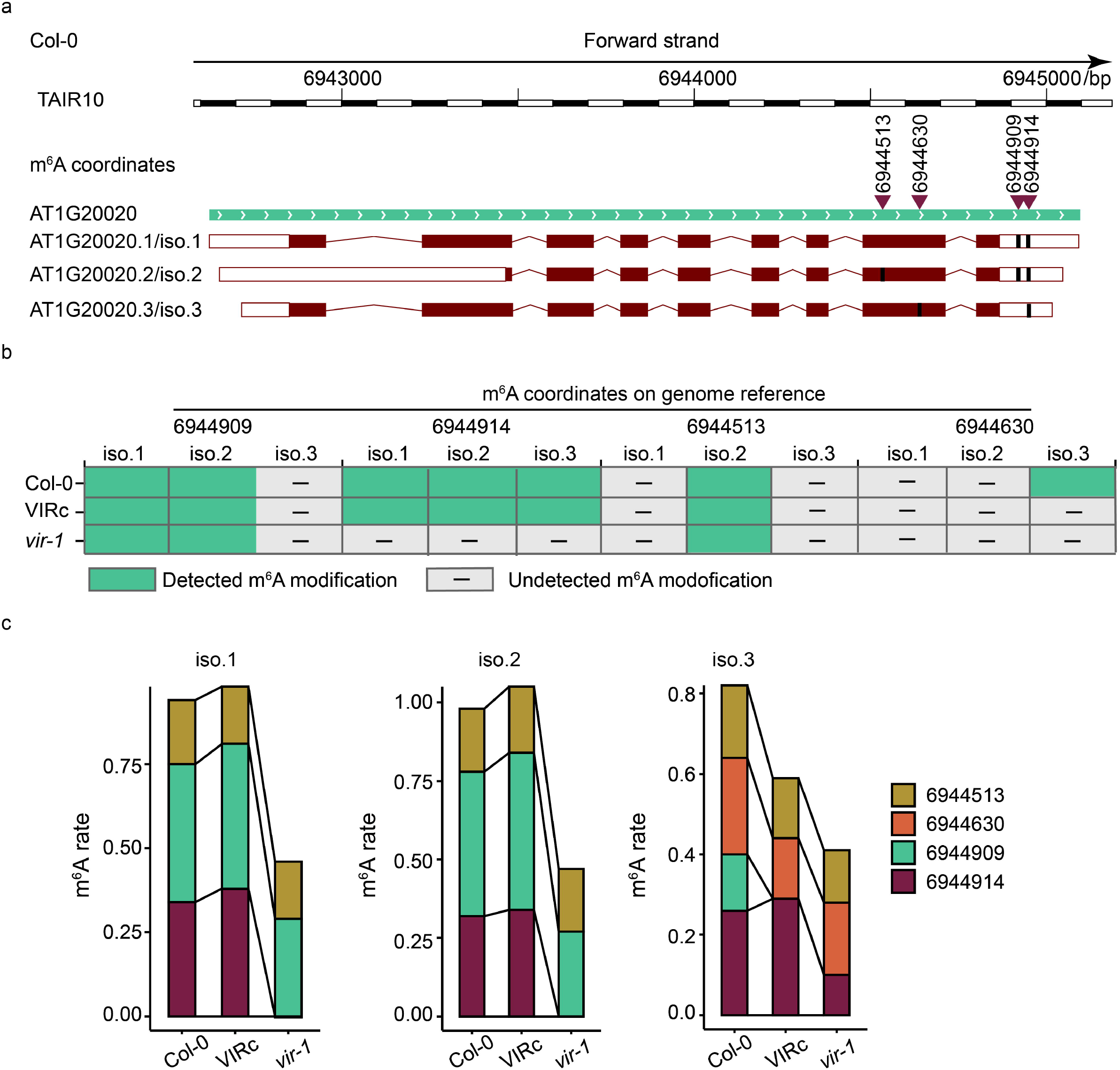
*DENA* identified m^6^A sites on different isoforms of single genes. (a) Cartoon displays three isoforms of AT1G20020. The different m^6^A sites on three isoforms are marked with dark bold lines, respectively. The m^6^A locations on the genome reference are indicated by red inverted triangles. (b) The distribution of seven high-confidence m^6^A sites on the three isoforms of AT1G20020 in Col-0, VIRc and *vir-1* lines, respectively. (c) The changes of m^6^A rate for the m^6^A sites in three isoforms of AT1G20020 from Col-0, VIRc and *vir-1* lines, respectively.

We next examined them in VIRc and *vir-1* lines, and six out of seven m^6^A sites in three isoforms were shared by both Col-0 and VIRc. However, only three sites were detected in *vir-1* (Figure 5b). For example, for the m^6^A site that was corresponding to the position 6560974 on the chromosome 1 of TAIR10, the modification rate was 0.59 and 0.42 in isoforms AT1G19000.1 and AT1G19000.2 of Col-0, respectively, which were consistent with the 0.518 and 0.406 in VIRc. In *vir-1*, however, this site was only identified in AT1G19000.2, with significant reduction of modification rate, only 0.15 (Figure 5c). Collectively, these results demonstrated the ability of *DENA* in m^6^A detection in different isoforms of single genes, which will be helpful for study the functions of m^6^A in RNA splicing.

### Profiling the m^6^A modification in *mtb* and *fip37-4 A.thaliana* mutants

We used *DENA* to quantify the m^6^A modification of three *A.thaliana* samples, Col-0, *mtb* and *fip37-4*, generated in this study (Additional file 2). We obtained 59827, 69504 and 55134 “RRACH” sites that were supported by at least 50 direct RNA-Seq reads in Col-0, *fip37-4*, and *mtb*, respectively. Among them, 19672 (32.88%), 18606 (26.77%), and 14068 (25.52%) m^6^A sites were identified in Col-0, *fip37-4*, and *mtb*, respectively, which showed a high overlap among them (Figure 6a). The preference for them to be present near the stop codon and within the 3’UTR in transcripts were confirmed (Figure 6b). We found the modification rates in both *fip37-4* and *mtb* were significantly decreased compared to that in Col-0, agreeing with the LC-MS/MS analysis (Figure 2a; Figure 6c; Additional file 1: Fig. S4a). In addition, the modification rates were highly correlative between *fip37-4* and *mtb* (Additional file 1: Fig. S4b).

**Figure 6.**
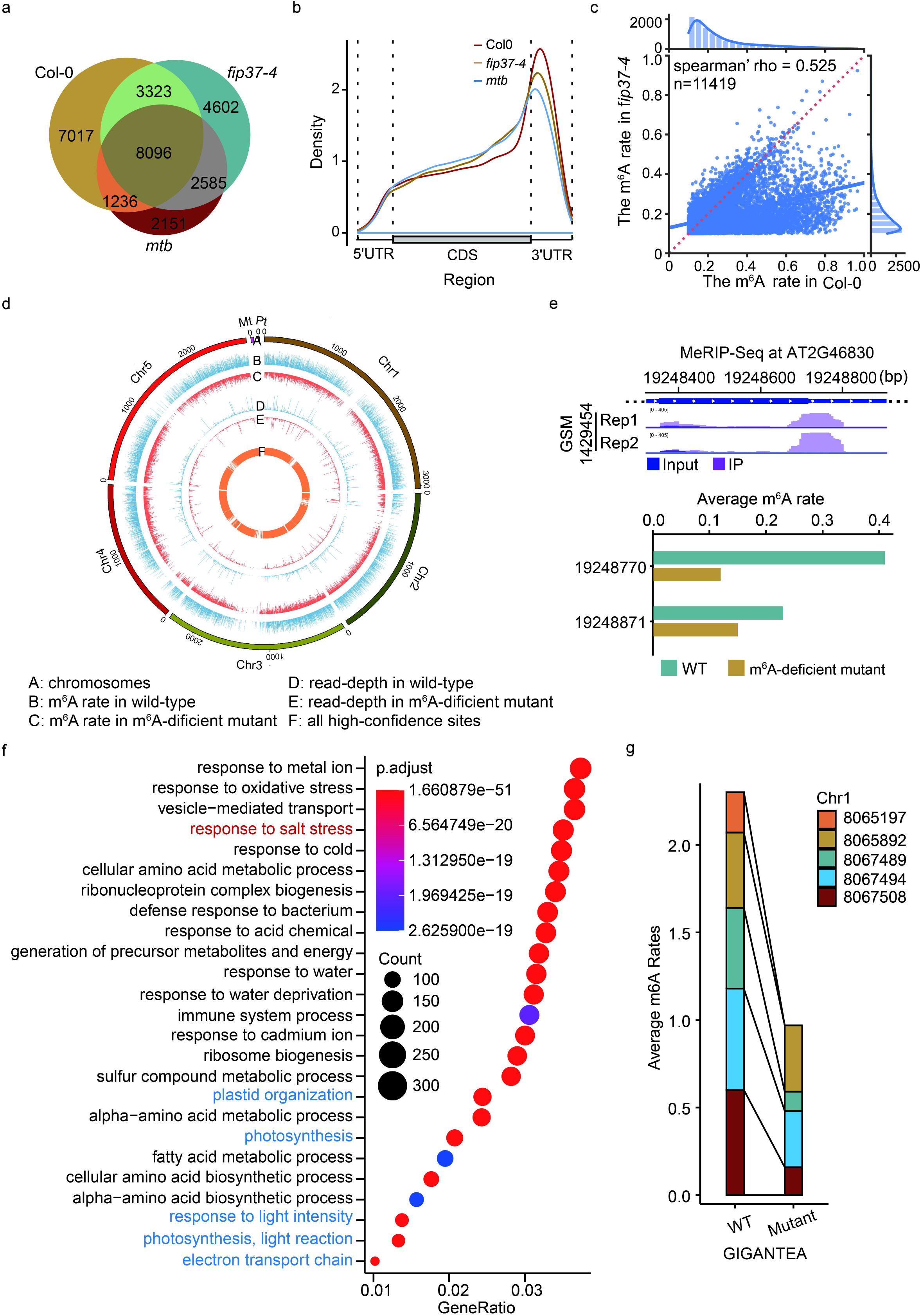
Profiling m^6^A modification in Col-0, *fip37-4*, and *mtb A.thaliana* lines. (a) Veen diagram shows the common m^6^A sites among Col-0, *fip37-4*, and *mtb* samples generated in this study. (b) The m^6^A distribution on transcripts in Col-0, *fip37-4*, and *mtb*, respectively. (c) Jointplot shows the correlation of m^6^A rate at 11419 overlapped sites between Col-0 and *fip37-4*. (d) Circos shows the profile, sequencing depth and modification rate of m^6^A sites in wild-type and m^6^A-deficient mutant with single-nucleotide resolution, respectively. (e) The top is the m^6^A peaks identified in CCA1 using MeRIP-seq in previous study, and the below is the average m^6^A rate of two high-confidence m^6^A sites identified by *DENA* from CCA1 gene in wild-type and m^6^A deficient mutant of *A.thaliana*, respectively. (f) Point maps the top 25 terms of Gene Ontology enrichment result with all m^6^A-modified genes. (g) The changes of average m^6^A rate at the five high-confidence m^6^A sites from GIGANTEA gene in wild-type and m^6^A-deficient mutant of *A.thaliana*, respectively.

To generate a richer resource of m^6^A modification with single-nucleotide resolution in *A.thaliana*, we combined all m^6^A sites identified by *DENA* in wild-type, VIRc and three mutants, *vir-1, fip37-4*, and *mtb*, and obtained 68136 m^6^A sites within 10223 genes in TAIR10 reference. A clearly reduced average modification rate was observed in the m^6^A-deficient mutant compared to wild-type (Figure 6d). For example, two high-confidence m^6^A sites at position 19248770 and 19248871 were identified by *DENA* in CCA1 (AT2G46830) of wild-type *A.thaliana*, which were identified by MeRIP in previous study[18]. However, the average rates were significantly decreased at both sites in the m^6^A-deficient mutant, especially at the position 19248770 (Figure 6e). In addition, Liu *et al*. established a m^6^A database, named REPIC[28], based on publicly available m^6^A-seq data. We compared the Arabidopsis m^6^A sites identified by DENA to the m^6^A data collected in the REPCI, and 48744 (71.54%) m^6^A sites from our model were covered by REPIC.

We performed the Gene Ontology analysis on all m^6^A-modified genes for functional insights into m^6^A modification in *A.thaliana*. These m^6^A-modified genes were significantly enriched in processes involved in the response to the abiotic and biotic stresses, biosynthetic process, and metabolism in biological process (Figure 6f). A recent study found that RNA m^6^A was important for salt stress tolerance in *A.thaliana*[29]. Consistently, a process of response to salt stress was enriched in our result. Additionally, photosynthesis, light reaction, and electron transport chain were also enriched, in agreement with the previous result that m^6^A modified transcripts were highly enriched in chloroplast/plastid and protein transport/localization categories[18]. For instance, GIGANTEA (AT1G22770), involved in phytochrome signaling[30] and salt response[29] in *A.thaliana*, was identified to contain m^6^A peaks in the 3’UTR[29]. Five high-confidence m^6^A sites were detected by *DENA* in its transcripts, including three in the 3’UTR and two in the 9^th^ exon. The modification rates of three sites from 3’UTR and one site from 9^th^ exon were significantly reduced in the m^6^A-deficient mutant (Figure 6g).

## Conclusions

m^6^A is the most abundant modification in mRNA and it is involved in many aspects of RNA functions. In this study, we developed a neural network, *DENA*, that can detect and quantify m^6^A in Nanopore direct RNA-Seq data at single-nucleotide resolution, providing a robust tool for transcriptome-wide profiling of m^6^A modification.

Using the second-generation sequencing, experimental approaches were developed to detect RNA m^6^A modification relying on immunoprecipitation[31-35], RNA editing via cytosine deaminase[36] or RNA digestion via m^6^A-sensitive enzyme[37, 38]. However, they require reverse transcription, specific antibodies or enzymes, and sophisticated experimental procedures, which may introduce biases and variations in their application.

Nanopore direct RNA-Seq can detect electrical signals of RNA m^6^A modification[32], which overcome these limitations relying on second-generation sequencing-related experimental approaches. One method identified m^6^A by comparing direct RNA-Seq data of wild-type with that of a matched m^6^A-deficient or hypomethylated sample[9]. However, the modification rate can not be accurately quantified, and the m^6^A-deficient or hypomethylated samples are difficult to generate, representing the major barriers in its application in profiling m^6^A modification. An alternative approach is to detect m^6^A modification based on their unique electrical signal fingerprints in direct RNA-Seq data. However, previous attempts were based on the data from *in vitro* synthetic RNAs in which all “A” residues are replaced by m^6^A, for training machine learning models[8, 14]. They suffered lower performance on m^6^A detection in *in vivo* transcripts as they are likely distorted due to superimposed electrical signals generated by clustered m^6^A residues in *in vitro* synthetic sequencing data. By contrast, *DENA* was developed with the direct RNA-Seq data from *in vivo* transcripts, and it can detect and quantify m^6^A at single-nucleotide resolution in the absence of m^6^A-deficient mutants.

The accuracy and reliability of *DENA* in m^6^A detection and quantification were validated in Arabidopsis and human data. On testing direct RNA-Seq data, *DENA* achieved higher accuracy of m^6^A identification (90%) in the m^6^A sites detected by miCLIP in *A.thaliana* (Figure 4). Analysis on m^6^A quantification for known sites in *Homo sapiens* showed that *DENA* had a higher correlation with those of the *SCARLET* method[22] than *Nanom6A* (Figure 4). *DENA’s* better performance may be due to the following reasons: First, *DENA* was trained on the direct RNA-Seq data containing the naturally occurring m^6^A sites that are different from the *in vitro* synthetic data with clustered m^6^A residues, eliminating the effect of superimposed signals in *in vitro* data (Figure 1). Second, a recent study found that the electrical signals of Nanopore sequencing displayed complex heterogeneity in methylation events[39]. Our *in vivo* training data extracted from over 3000 m^6^A sites, covered diverse combinations of sequence context and overcame the limited diversity of the 130 sites containing “RRACH” motif in the *in vitro* synthetic data. As proposed in previous studies, the performance of m^6^A detection was improved on the sophisticated neural network, like *DENA*, by training with our extended *in vivo* transcribed data.

The inability to identify m^6^A sites specific to different isoforms of gene transcripts hampered functional studies of m^6^A in RNA splicing[4]. Another important improvement of *DENA* is that it can identify and quantify m^6^A modification on different isoforms of gene transcripts (Figure 5), which is unavailable in previously methods. *DENA* assigns sequencing reads to transcript isoforms, which can obtain the isoform compositions and detect the modification sites associated with isoforms. In addition, we evaluated the m^6^A profiles of *fip37-4* and *mtb* mutants at single-nucleotide resolution using *DENA*, and found a high correlation between the two mutant lines (Additional file 1: Fig. S4b). The results suggested overlapping function between FIP37 and MTB as they are both subunits of the *N*^6^-methyltransferase complex catalyzing the formation of m^6^A modification in RNA.

In summary, *DENA* (available at https://github.com/weir12/DENA) is the first attempt to use direct RNA-Seq data of *in vivo* transcribed mRNAs to train neural network model for m^6^A detection, which overcame the weakness of previous methods trained using *in vitro* synthetic RNAs with all adenosine residues replaced by m^6^A. *DENA* shows a better performance than previous methods and is robust in handing direct RNA-Seq data cross different species. Moreover, we noticed that there were few reads of long non-coding RNAs (lncRNAs) in the human data of direct RNA sequencing. We speculated that most lncRNAs were left out due to lacking poly(A) tail when purifying mRNA by poly(T). Thus, we suggest it may be advantageous to use the poly(A) polymerase to add poly(A) tails to the 3’-end of lncRNAs for detection of m^6^A modification on lncRNAs, before building the library for direct RNA sequencing. In addition, although our current work was developed using the R9.4 flowcells to perform direct RNA sequencing for m^6^A detection, our computational framework may be applied to new platforms of direct RNA sequencing technology, which keeps the utility of DENA in pace with future technology development. Collectively, our study provides a useful framework for development of novel models in the detection of other types of RNA modifications using Nanopore direct RNA sequencing.

## Methods

### Plant materials and growth conditions

All *A.thaliana* lines were derived from the Columbia (Col-0) accession. Briefly, the hypomorphic allele of FIP37 was generated by a T-DNA insertion within its 7^th^ intron (*fip37-4* SALK_018636 allele, termed *fip37-4*) as reported[17]. The MTB-deficient mutant was generated by rescuing the null mutation through the embryo-specific expression of MTB driven by the ABI3 promoter (*ABI3prom:MTB* complemented *mtb* WiscDsLox336H07 allele, termed *mtb*), because null mutations in MTB are embryonic lethal. Seeds that were stratified at 4°C for two days, were sown on MS medium plates, before they were cultivated in a controlled environment at 22°C under a 16h:8h photoperiod. Seedlings were harvested 14 days after transfer to 22°C. All materials were frozen immediately after harvest and stored for standby at -80°C.

### RNA isolation

RNAprep pure Plant Kit (TIANGEN BIOTECH CO., LTD) was used to extract total RNA from all samples and Oligotex mRNA Mini Kit was used to purify the polyadenylated mRNA from total RNA. All mRNAs were frozen immediately and were stored at -80°C.

### LC-MS/MS

LC-MS/MS was performed as described[40]. Briefly, about 500□ng mRNA of each sample was digested by 2 U nuclease P1 (NEB) in 4 μL of 10X Nuclease P1 Reaction Buffer at 37°C for 30 min, followed by additions of 1 U Thermosensitive Alkaline Phosphatase (1□U/μL) and Thermo Scientific™ FastAP™ reaction buffer. The final mixture incubated at 37□°C for 2□h. The sample was dissolved in 50 μl anhydrous methanol after drying under vacuum, and 10 μL of the solution was used for LC-MS/MS analysis. The samples, and m^6^A and adenosine (A) standards were separated by reverse-phase ultra-performance liquid chromatography on a C18 column with online mass spectrometry detection using Agilent 6500 QQQ triple-quadrupole LC mass spectrometer in positive electrospray ionization mode. The nucleotides were quantified by using the nucleotide-to-base ion mass transitions of *m/z* 282.0 to 150.1 (m^6^A), and *m/z* 268.0 to 136.0 (A). Three biological replicates for each strain (Col0, *mtb*, and *fip37-4*) were performed and the LC-MS/MS analyses were performed simultaneously on the same machine.

### Nanopore direct RNA-Seq and data processing

Nanopore direct RNA sequencing was performed using MinION MkIb with R9.4 flowcells at Nextomics Biosciences Co., Ltd (Wuhan, China). Briefly, mRNA was isolated from about 75 μg total RNA for each sample using the Dynabeads mRNA purification kit (Thermo Fisher Scientific). The quality and quantity of mRNA were assessed using the NanoDrop 2000 spectrophotometer (Thermo Fisher Scientific). Then the nanopore libraries were prepared from 1 μg poly(A)+ RNA with the direct RNA sequencing Kit (SQK-RNA001, Oxford Nanopore Technologies) according to the manufacturer’s instructions. The adapter of poly(T) was ligated to the mRNA using T4 DNA ligase (New England Biolabs) in the reaction buffer for 10 min at 25□. And then the cDNA was synthesized using SuperScript III Reverse Transcriptase (Thermo Fisher Scientific) with the oligo(dT) adapter and incubate at 50° C for 50 min, then 70° C for 10 min, and bring the sample to 4° C before proceeding to the next step. Then the RNA-DNA hybrid was purified by Agencourt RNAClean XP magnetic beads (Beckman Coulter). The sequencing adapter was ligated to the mRNA from RNA-DNA hybrid using T4 DNA ligase (New England Biolabs) in the NEBNext Quick Ligation Reaction Buffer (New England Biolabs) for 10 min at 25°C followed by a second purification step using Agencourt beads (as described above). 1 μl reverse-transcribed and adapted RNA were quantified by the Qubit fluorometer DNA HS assay. Then libraries were loaded on GridION using R9.4 flowcells (Oxford Nanopore Technologies) with Library Loading Bead Kit (EXP-LLB001, Oxford Nanopore Technologies) and sequenced using a 48-h run time. Three biological replicates for each strain were sequenced in independent machines and days.

Base-calling was performed with *Guppy* (version 2.3.4) using default parameters. Reads were aligned to the TAIR10 genome and Araport11 transcriptome references of *A.thaliana* using *minimap2* tool (version 2.813)[41]. Sequence Alignment/Map (SAM) and BAM file manipulations were performed using *samtools* (version 1.956). The length of direct RNA-Seq reads was calculated using the scripts of *wub* tool (Oxford Nanopore Technologies Ltd).

### Detecting m^6^A sites in the m^6^A-dificient mutants using *differr*

All fast5 files from direct RNA-Seq was bse-called to fastq files by *Guppy* (version 3.2.4) in this study. The sequence data (fastq files) from m^6^A-dificient mutants and wild-type *A.thaliana* were aligned to TAIR10 reference using *minimap2*. The differential sites between mutant and wild-type using was identified using *differr*[9] with default parameters. The genomic coordinates were converted to corresponding coordinates of transcriptional isoforms using the R package *GenomicFeatures* (v1.40.0)[42]. The relative distance is obtained by subtracting the coordinate of the differential sites from that of the core “A” base in the nearest m^6^A motif “RRACH”. The center “A” residue of “RRACH” that containing differential “error” sites within 5 nucleotides, were regarded as m^6^A modification sites. The electrical signals of direct RNA-Seq data were extracted using *Tombo* tool[43].

### Extracting native training data from direct RNA-Seq reads of *A.thaliana*

All fast5 files of VIRc and *vir-1* samples was base-called to fastq reads by *Guppy* (version 3.2.4) and subsequently fastq reads were aligned to Araport11 transcriptome reference. We used *Tombo* re-squiggle algorithm to correct the raw base-calling sequence based on an expected electric level model, and then assigned the corrected base to the raw electric signal segment. After correction, we decided to extract the m^6^A-modified and unmodified training data from the direct RNA-Seq reads of VIRc and *vir-1 A.thaliana* lines, respectively. For 3106 common m^6^A sites identified by *differr* tool across C*m*, C*f* and V*v*, we regarded the 11 nucleotides of each site (the nucleotide of m^6^A site and five nucleotides upstream and downstream) as a “region”, and obtained 3106 “region”. We then extracted the fragments from all base-calling reads at each “region” in VIRc and *vir-1*, respectively. If a fragment from VIRc sample occurred mismatches, we labeled this fragment as a positive event, and extracted its matrix of features (mean, median, standard deviation, dwell time, and base quality) from nanopore electric signals within the “RRACH” window. Conversely, if one fragment from *vir-1* mutant did not occur mismatches, we labeled this fragment as negative event, and extracted its matrix of features as above. The dataset containing all matrices of positive events is called m^6^A-set, while the dataset including all matrices of negative events is called A-set. In all, we obtained the 2251835 matrices of features (mean, median, standard deviation, dwell time, and base quality) from the events of direct RNA-Seq reads, which contains 507066 in m^6^A-set and 1744769 in A-set.

### Training neural network model with the m6A-set and A-set

Bidirectional Long Short-Term Memory (Bi-LSTM) neural network is a variant of Recurrent neural network (RNN)[44]. It can solve the long-term dependency problem (gradient exploding and gradient vanishing) of general RNN, and has achieved superior performance in Time Series Prediction, Natural Language Processing, and Machine Translation []. Bi-LSTM can capture the rules in the time-series data, such as direct RNA-Seq data, and has been used to detect DNA base modification on Oxford Nanopore sequencing data[15, 16]. Therefore, we consider that Bi-LSTM can be designed to detect of m^6^A methylation based on nanopore direct RNA sequencing. Then, we divided these labeled features into 12 consensus patterns (“AAACA”, “AAACC”, “AAACT”, “AGACA”, “AGACC”, “AGACT”, “GAACA”, “GAACC”, “GAACT”, “GGACA”, “GGACC” and “GGACT”) according to the sequence. For each pattern, we further divided its features into training and testing datasets at a ratio of 7:3 for training independent sub-model using Bi-LSTM. In our study, the Bi-LSTM is constructed with three hidden layers after hyperparametric optimization, where bidirectional RNN considers both forward and reverse data flow from neighborhood bases. Each LSTM unit contains multiple hidden nodes. In brief, the flow of tensors during forwarding and backpropagation is visualized in Figure 3, and the output layer of the Bi-LSTM is flattened and feed into a three-layer fully connected neural network for obtaining the predicted results and calculating the loss function. Samples from different reference sites and reads were randomly shuffled during the training. The prediction labels are scaled by the SoftMax function, and cross-entropy would be minimized to tune the parameters. We use an Adam optimizer with an initial learning rate of 0.0005 to update the weight and bias. However, with the increase of epochs, the learning rate will be reduced. Thus, we decay the learning rate of each parameter group by gamma (0.1) every epoch. To avoid over-fitting, we stop training and retain the model with the best verification set, if three successive epochs do not improve the accuracy in the verification set. The model containing 12 sub-models is the finally available neural network model named *DENA*. The implementation of *DENA* relies on the PyTorch, a python-based computing package. The NVIDIA Tesla V100 was used to speed up the both training and testing of our model.

We used following parameters for evaluating our model:

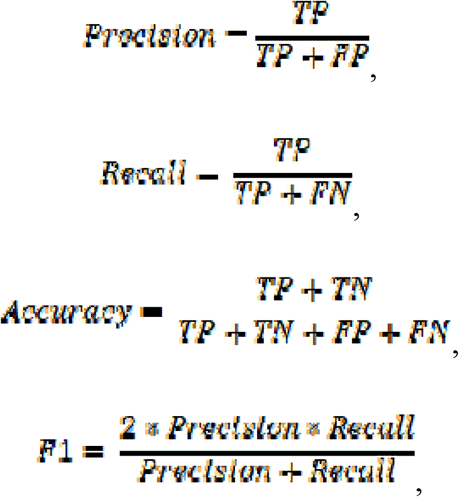

where *TP, FP, TN, FN, F1* are true positives, false positives, true negatives, false negatives and F1 score, respectively. We also measure a predictive power of *DENA* using the area under the curve (AUC) and the receiver operating characteristic curve (ROC)[45]. For *DENA* model, the input dataset was raw signals from Nanopore direct RNA sequencing. One output result was the predicted modification probability at each “RRACH” site on each nanopore signal, and another output result was the m6A modification rate at each “RRACH” site on reference sequence. The modification rate of each “RRACH” site was calculated as follows:

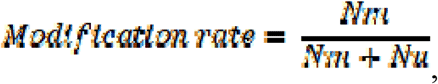

where *Nm* and *Nu* represent the number of signals with a predicted modification probability of not less than 0.5 and the number of signals with a predicted modification probability of less than 0.5 at each site, respectively

### Identification and quantification of m^6^A modification in *A.thaliana* and human using *DENA*

After aligning fastq reads to transcriptome reference, the raw signals are assigned to the reference based on an expected electric level model by re-squiggle algorithm of *Tombo* tool. *DENA* firstly calculates the position of all “RRACH” motifs in the transcriptome reference, and the following extracts the features from the electric signal. Finally, *DENA* calculates the depth of sequencing at each position and performs the prediction of both transcriptional coordinate and modification rate (m^6^A reads/total reads) of m^6^A from all “RRACH” sites. The sequencing depth of each sample could limit the number of genes and positions that can be identified because lowly expressed genes being potentially excluded. To maximize the coverage depth of genes and improve the reliability of the m^6^A ratio, we pool all reads across all replicates of each sample of *A.thaliana*. For the direct RNA-Seq data of human cells that was established by Ploy N. Pratanwanich *et. al*., we pool all reads across replicate1 (rep1) and replicate3 (rep3) of wild-type, in order to keep the same data size of replicate1 (rep1) of METTL3-knockout mutant cells. To further reduce the impact of sequencing depth and effectively control the false positives, we filtered sites with the following steps: (I) supported by at least 50 reads coverage; (II) the site with modification rate not less than 0.1 as m^6^A sites; (III) the site with modification rate greater than 0.2 as high-confidence site; (IV) the site with m^6^A rate between 0.1 and 0.2 as low-confidence site. The coordinate of m^6^A sites on transcriptome reference was converted to corresponding coordinate on genome reference by in-house python scripts. The consensus motif is plotted by the *MEME* tool[46] and the distribution of m^6^A on transcripts is obtained using *Guitar* package (version 2.8.0)[47]. The m^6^A profiles of *A.thalian* are plotted by *Circos* software (version 0.69-9). The m^6^A sites in the profiles is the union of the sites identified by *DENA* in VIRc, Col-0, and three m^6^A-deficient mutants of *A.thaliana*. The average modification rate of m^6^A site in wild-type *A.thaliana* was the mean of rates from VIRc and Col-0 samples. The average modification rate of m^6^A site in m^6^A-deficient mutant was the mean of rates among three mutants, *vir-1, fip37* and *mtb*. The GRCh38.p13 reference is used for human, and Araport11 and TAIR10 references for *A.thaliana*. All transcriptome and genome references are downloaded from Ensembl database.

### Identification of *DENA*-predicted m^6^A modification in *A.thaliana* using experimental method

We confirmed the DENA-predicted m^6^A sites in Arabidopsis using *SELECT* method described by Xiao et al.[24]. As shown in Figure 4h, one synthetic DNA probe with PCR adapter (named Dp) and another synthetic DNA probe (named Up) complementarily anneal to RNA but leave a gap of three nucleotides opposite to the candidate site to be identified. As described in other work[26], most mammalian m^6^A sites are within the G(A/m^6^A)CU motif. To minimize the over extension of the Up probe by *Bst* 2.0 DNA polymerase, only dATP, dTTP, dGTP, but not dCTP are added to prevent extension across guanosine in G(A/m^6^A)CH (H=A, C or U) [26]. If a m^6^A modification present in this candidate site of RNA template, it will selectively hinder *Bst* 2.0 DNA polymerase mediated single-base elongation of the Up probe and stop it at the “C” base behind m^6^A-modified base. However, if non-modification presents in candidate site, the elongation will continue to the candidate “A” base behind the “G” base. Note although this step is not 100% efficient (a small number of elongation products will still reach the m^6^A-modified site), *SplintR* ligase will be block their connection with Down probes due to the existence of m^6^A modification at candidate site in the next ligation step. Meanwhile, we select an unmodified “A” base nearby candidate m^6^A-modified site as the control. By comparing the cycles of qPCR between the candidate site and the control unmodified “A” base, we can confirm whether there is m^6^A modification at the candidate site.

In detail, the 5’ phosphorylation of down DNA probe for subsequent ligation was produced by T4 Polynucleotide Kinase within a mixture containing 15 U T4 Polynucleotide Kinase (Vazyme Biotech Co. Ltd.), 50 μM up DNA probe, 0.5 mM ATP and 1x T4 PNK buffer, at 37 °C for 30 min. Annealing of up and down DNA probes to RNA templates was conducted within the 15 μ L mixture containing 1.5 μ g total RNAs, 5 pmol 5’ phosphorylation of down DNA probe, 5 pmol up DNA probe and 0.2 μL RNase inhibitor. The mixture was incubated at 90 °C for 1 min, and then the temperature was reduced at an interval of 10 °C, with 1 min of incubation at each interval end point. After the temperature had dropped to 50 °C, the mixture was kept at 4 °C. Subsequently, the elongation reaction mixture consists of above 15 μL annealed mixture and another 50 μL mixture containing 50 nmol dDTPs (dATP, dGTP and dTTP) [26], 125 nmol ATP, 0.8 U *Bst* 2.0 DNA polymerase (NEB) and 0.2 μL RNase inhibitor (ThermoFisher Scientific) with 1x CutSmart buffer (NEB). The 65 μL mixture for elongation was incubated at 45 °C for 30 min. Finally, a 100 μL ligation reaction mixture consisted of above 65 μL elongation mixture and 35 μL another mixture containing 8 U *SplintR* ligase (NEB), 20 nmol ATP, 25 μ L 50% PEG8000 (RNase-free, Beyotime Biotechnology) and 1x CutSmart buffer. The final mixture was incubated at 37 °C for 40 min, and then was kept at 4 °C. The ligation product was diluted for 100-folds, and then add 1 μL into a 20 μL scale qPCR reaction with Taq Pro Universal SYBR qPCR Master Mix (Vazyme). The qPCR was performed on QuantStudio 3 Real-Time PCR System with the following parameters: 94 °C, 30s; (95°C, 5s; 57 °C, 20s; 72 °C, 10s) × 45 cycles; 4°C, hold. Data was analyzed using Design and Analysis Setup software (version 2.6). All DNA probes used in this study were list in Additional file 1: Table S4.

### Availability of data and materials

All accessions of datasets used in our study were list in the Additional file 3. The direct RNA-Seq reads of wild-type, *vir-1* and VIR-complemented (VIR::GFP-VIR) *A.thaliana* lines are downloaded from the European Nucleotide Archive (ENA) under accession PRJEB32782[9]. The direct RNA-Seq reads of wild-type and METTL3 knock-out *H. sapiens* cells are downloaded from the ENA under accession PRJEB40872[23]. The direct RNA-Seq reads of *in vitro* synthetic transcripts are downloaded from the ENA under accession PRJNA511582[8]. All direct RNA-Seq reads of wild-type, *fip37-4* and *mtb A.thaliana* lines generated by this study have been submitted to the ENA under accession PRJEB45935[48], and National Genomics Data Center, China National Center for Bioinformation (CNCB-NGDC) under project accession PRJCA007105 and GSA accession CRA005317[49]. All in-house python scripts used in this study are publicly available as part of *DENA* that can be download and used from Github with MIT License (https://github.com/weir12/DENA)[50], and zenodo (https://zenodo.org/record/5603381)[51].

## Acknowledgements

We thank the Nextomics Biosciences Co., Ltd for performing Nanopore direct RNA sequencing, and thank Professor Lianfeng Gu for his help in the use of *Nanom6A* software.

## Author contributions

X.L., P.H. and J.W. conceived this project and directed the study. L.C., and J.G. assisted H.Q. to perform the experiments. L.O. extracted training data sets and trained the model. H.Q. analyzed data and prepared the manuscript.

## Ethics approval and consent to participate

Not applicable.

## Consent for publication

Not applicable.

## Supplementary Information

**Additional file 1:**

**Fig. S1** Nanopore direct RNA-Seq implementation and m^6^A detection with *differr* tool.

**Fig. S2** Training DENA.

**Fig. S3** Confirming the reliability of *DENA* in m^6^A quantification.

**Fig. S4** The correlation of modification rate between wild-type and m^6^A-deficient *A.thaliana* mutant.

**Table S1** Sequencing statistics of poly(A) selected RNAs in biological triplicates from Col0, *mtb*, and *fip37-4* using direct RNA-Seq, respectively.

**Table S2** The performance of the *DENA* prediction model that was evaluated with metrics including accuracy, recall, precision, F1-score.

**Table S3** The comparison of m^6^A modification rates between *DENA* and other methods (containing *xPore, Nanom6A, SCARLET, LEAD-m6A-seq* and *Deoxyribozyme*-based Method) at the previously identified m^6^A sites in human. NT: Not detected; -: Not identified; Y: identified as m^6^A site.

**Table S4** DNA probes used in the SELECT assay.

**Additional file 2** *DENA* output containing m^6^A sites in Col-0, *fip37-4* and *mtb*, and all nonredundant m^6^A sites identified across all Arabidopsis lines used in this study.

**Additional file 3** List of datasets used in this study.

