## Additional file 1 for "DENA: training an authentic neural network model using Nanopore sequencing data of Arabidopsis transcripts for detection and quantification of *N*^6^-methyladenosine on RNA"

### Contents

**Supplementary Figures**

**Supplementary Tables**

### Supplementary Figures

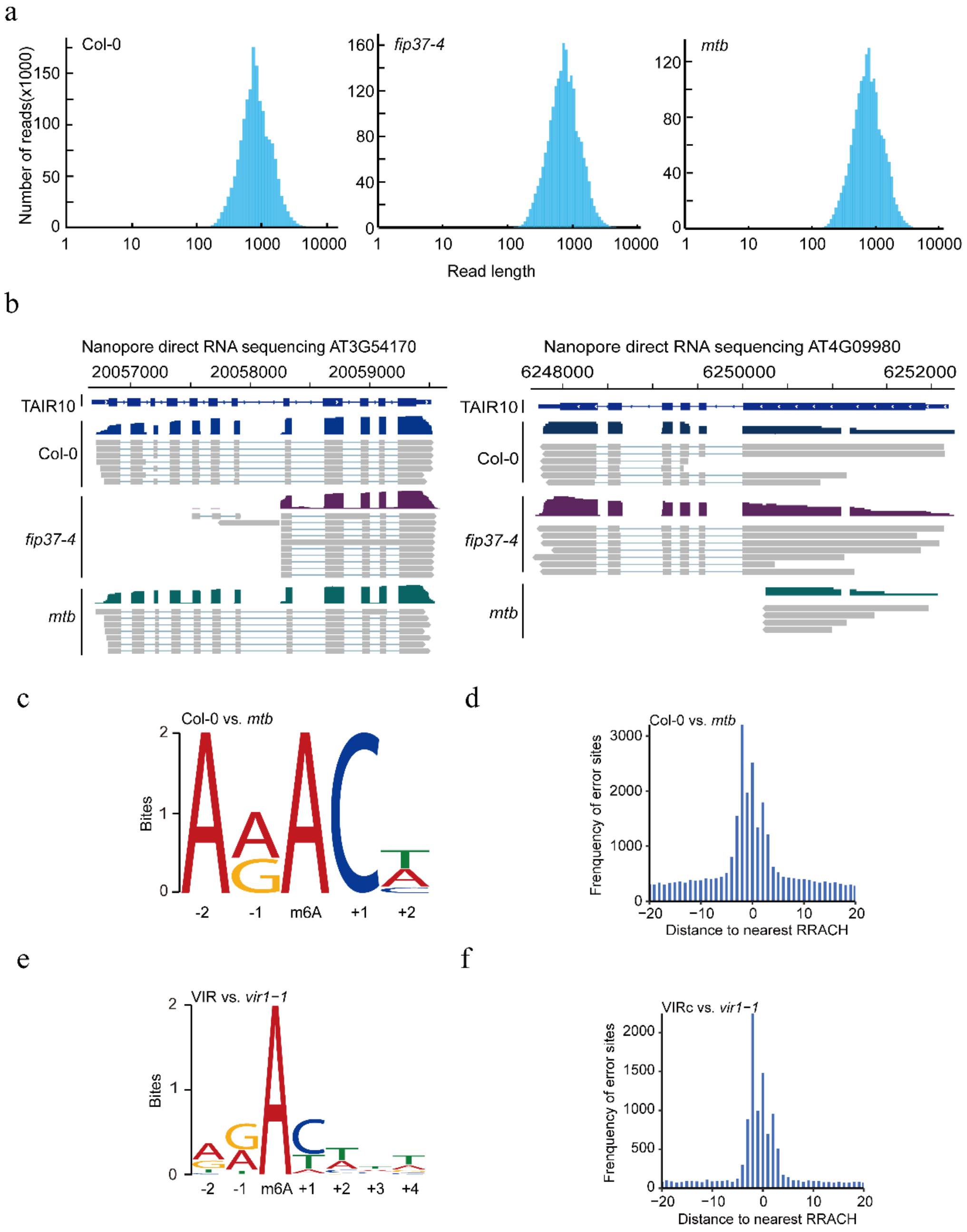


**Fig. S1 Nanopore direct RNA-Seq implementation and m$^6$A detection with *differr* tool**. (a) The distribution of direct RNA-Seq reads from Col-0, *fip37-4*, and *mtb*, respectively. (b) IGV shows the alignment of direct RNA-Seq reads for FIP37 (AT3G54170) and MTB (AT4G09980) gene in Col-0, *fip37-4*, and *mtb*, respectively. (c) and (e) are the consensus motifs in C*m* and V*v*, respectively. (d) and (f) are the distribution of distances between the differential sites and its nearest “RRACH” sequences in C*m* and V*v*, respectively.

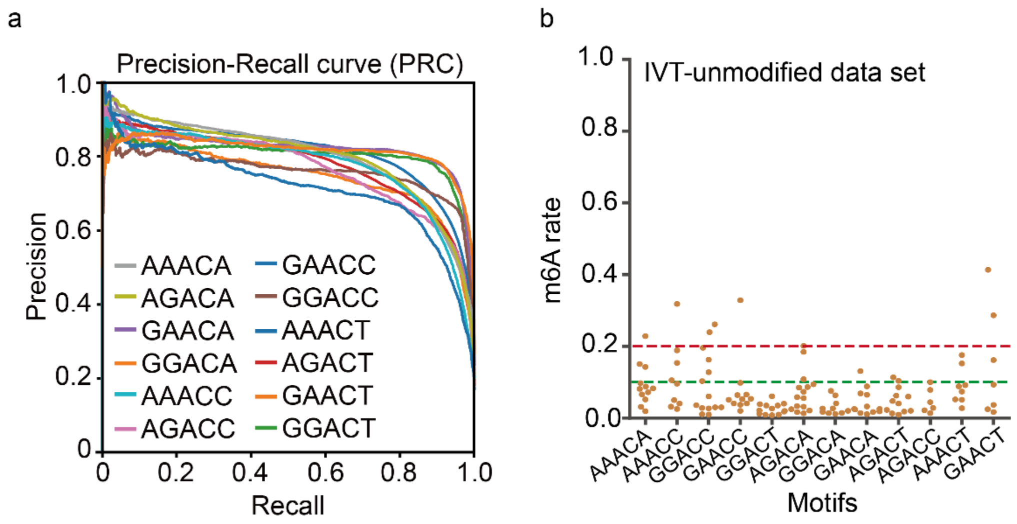


**Fig. S2 Training DENA.** (a) The Precision-Recall curve (PRC) of *DENA*. (b) The $m^6A$ prediction for unmodified training data from *in vitro* synthetic RNAs using *DENA*. The green and red lines show the $m^6A$ rate with 0.1 and 0.2, respectively.

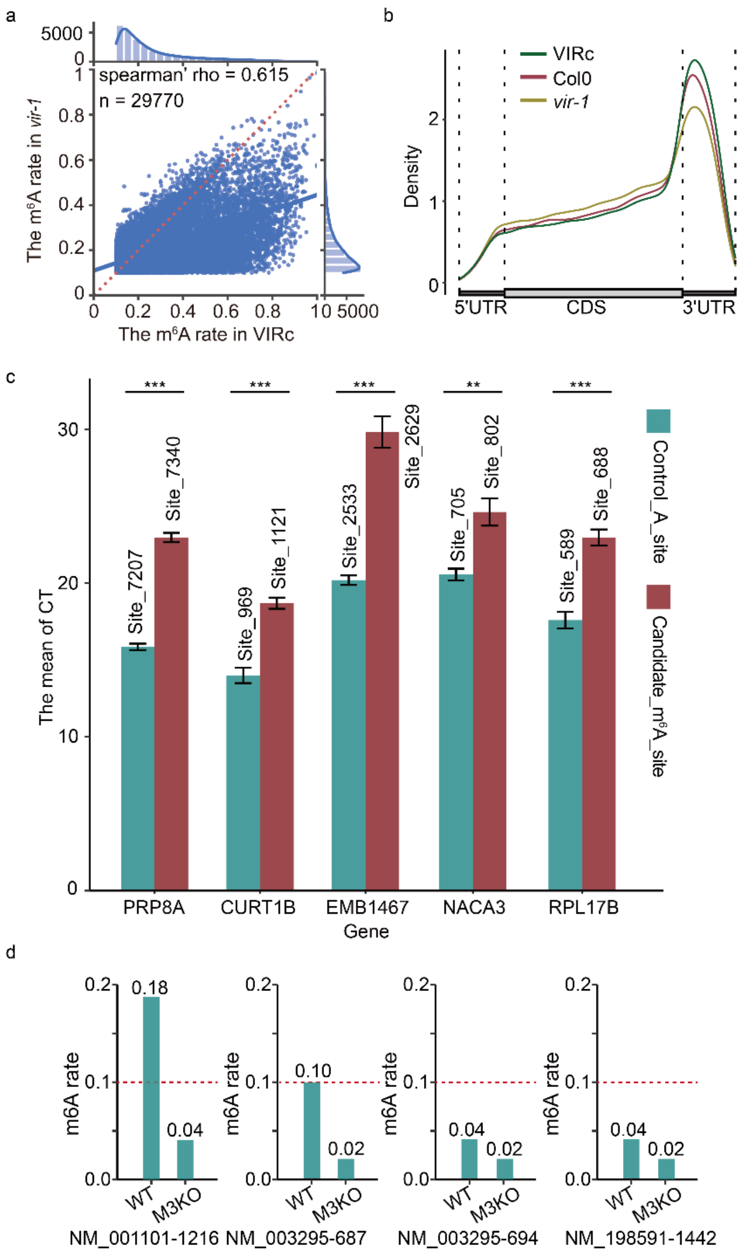


**Fig. S3 Confirming the reliability of *DENA* in m$^6$A quantification.** (a) Jointplot shows the correlation of m$^6$A rates from 29770 intersected sites between *vir-1* and VIRc. (b) The m$^6$A distribution on transcripts in Col-0, VIRc and *vir-1*, respectively. (c) The identification of five m$^6$A sites predicted by DENA using qPCR (real-time quantitative PCR). Green shows the “A” base of control. Red shows the DENA-predicted m$^6$A site. P values from t-test (two-tailed) are shown on top of the bar plots. *p < 0.05; **p < 0.01; ***p < 0.001. (d) Bar plot shows the m$^6$A rates identified by *DENA* at NM_001101-1216, NM_003295-687, NM_003295-694 and NM_198591-1442 sites in WT and M3KO cells, respectively.

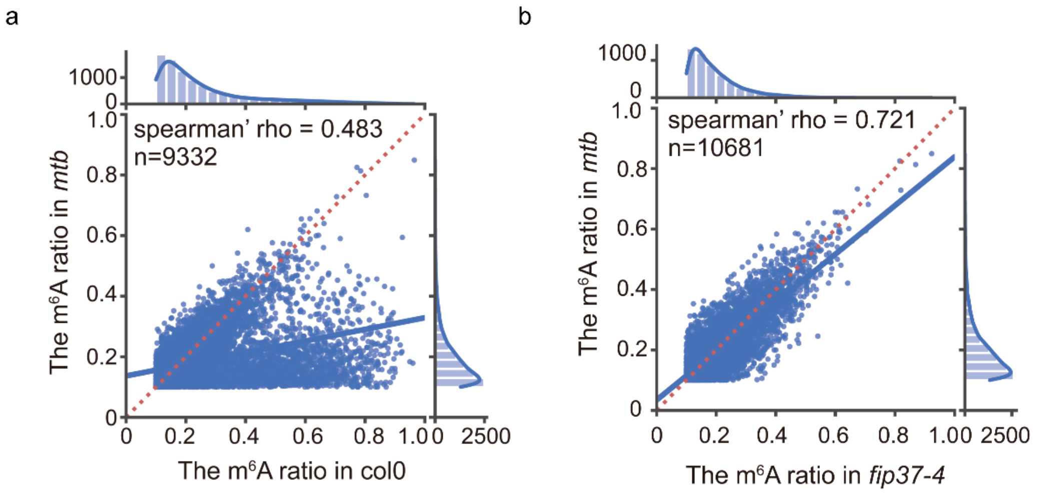


**Fig. S4 The correlation of modification rate between wild-type and m$^6$A-deficient *A.thaliana* mutant.** (a) Jointplot shows the correlation of m$^6$A rates at 9330 intersected sites between Col-0 and *mtb*. (b) Jointplot shows the correlation of m$^6$A rates at 29770 intersected sites between *fip37-4* and *mtb*.

**Supplementary Tables**

**Table S1** Sequencing statistics of poly(A) selected RNAs in biological triplicates from Col0, *mtb*, and *fip37-4* using direct RNA-Seq, respectively.

| | | Col0 | | | *fip37-4* | | | *mtb* | | |
|---|---|---|---|---|---|---|---|---|---|---|
| | | Col0_1 | Col0_2 | Col0_3 | *fip37_1* | *fip37_2* | *fip37_3* | *mtb_1* | *mtb_2* | *mtb_3* |
| Raw data | Number of reads | 355285 | 518005 | 1016370 | 621552 | 737486 | 1019317 | 380011 | 760082 | 758060 |
| | Mean read length | 1024.9 | 1044.1 | 740.1 | 891.5 | 906.9 | 682.7 | 916.5 | 913.3 | 731.2 |
| | Mean read quality | 7.8 | 7.8 | 7.7 | 8 | 7.9 | 7.6 | 7.9 | 8 | 7.7 |
| | Median read length | 859 | 874 | 638 | 774 | 785 | 577 | 787 | 786 | 616 |
| | Median read quality | 7.8 | 7.7 | 8 | 8 | 7.9 | 7.9 | 7.9 | 8 | 8 |
| | Read length N50 | 1162 | 1191 | 895 | 1011 | 1024 | 883 | 1032 | 1034 | 928 |
| | Total reads | 1889660 | | | 2378355 | | | 1898153 | | |
| | Total aligned reads | 1739015 | | | 2181217 | | | 1756890 | | |
| Quanlity Control (read-length >100bp and Q>7) | Number of reads | 352600 | 513852 | 841035 | 617154 | 731679 | 810171 | 377050 | 754448 | 622851 |
| | Mean read length: | 1027.7 | 1046.8 | 806.9 | 895 | 910.7 | 775.9 | 920.1 | 917 | 812.6 |
| | Mean read quality | 7.8 | 7.8 | 8.1 | 8 | 7.9 | 8 | 7.9 | 8 | 8.1 |
| | Median read length | 861 | 876 | 698 | 776 | 787 | 663 | 789 | 788 | 692 |
| | Median read quality | 7.8 | 7.7 | 8.1 | 8 | 7.9 | 8 | 7.9 | 8 | 8.1 |
| | Read length N50 | 1164 | 1193 | 918 | 1011 | 1024 | 918 | 1033 | 1035 | 962 |
| | Total reads | 1707487 | | | 2159004 | | | 1754349 | | |
| | Total aligned reads | 1687722 | | | 2117431 | | | 1714419 | | |

**Table S2** The performance of the *DENA* prediction model that was evaluated with metrics including accuracy, recall, precision, F1-score.

| **Motifs** | **Accuracy** | **Precision** | **Recall** | **F1-score** |
|---|---|---|---|---|
| AAACT | 0.8738 | 0.7044 | 0.8782 | 0.7818 |
| GAACT | 0.8832 | 0.6204 | 0.9113 | 0.7382 |
| GGACT | 0.9182 | 0.7705 | 0.8978 | 0.8293 |
| AGACT | 0.8798 | 0.6100 | 0.9161 | 0.7324 |
| GAACA | 0.9264 | 0.7341 | 0.9510 | 0.8286 |
| GGACC | 0.9191 | 0.6912 | 0.9059 | 0.7841 |
| AGACC | 0.8563 | 0.6005 | 0.9078 | 0.7228 |
| AAACA | 0.8396 | 0.6908 | 0.8396 | 0.7579 |
| AGACA | 0.8706 | 0.6161 | 0.9047 | 0.7330 |
| AAACC | 0.8835 | 0.6574 | 0.8678 | 0.7481 |
| GAACC | 0.8931 | 0.6133 | 0.8588 | 0.7156 |
| GGACA | 0.9142 | 0.7657 | 0.9257 | 0.8381 |

**Table S3** The comparison of m$^6$A modification rates between *DENA* and other methods (containing *xPore*, *Nanom6A*, *SCARLET*, *LEAD-m6A- seq* and *Deoxyribozyme*-based Method) at the previously identified m$^6$A sites in human. NT: Not detected; -: Not identified; Y: identified as m$^6$A site.

| gene | genoLoci | isoforms | transLoci | motif | *xPore* | | *Nanom6A* | | *DENA* | | *SCARLET* | *LEAD-m6A-seq* | *Deoxyribozyme* |
|---|---|---|---|---|---|---|---|---|---|---|---|---|---|
| | | | | | rate | | rate | | rate | | rate | rate | |
| | | | | | **WT** | M3KO | **WT** | M3KO | **WT** | M3KO | **WT** | **WT** | **WT** |
| ACTB | 5527743 | NM_001101.5 | 1217 | GGACT | **0.75** | 0.18 | **0.64** | 0.13 | **0.18** | 0.04 | **0.21** | **0.29** | **Y** |
| BSG | 583239 | NM_001322243.2 | 1340 | GGACT | **NT** | NT | **0.67** | 0.55 | **0.29** | 0.09 | **0.06** | **0.55** | **-** |
| | | NM_198589.3 | 1344 | GGACT | **1** | 0.24 | | | **0.36** | 0.09 | | | |
| | | NM_198591.4 | 1378 | GGACT | **1** | 0.19 | | | **0.16** | 0.02 | | | |
| | 583346 | NM_001322243.2 | 1447 | GAACT | **NT** | NT | **0.18** | 0.17 | **0.06** | 0.02 | **0.01** | **-** | **-** |
| | | NM_198589.3 | 1451 | GAACT | **NT** | NT | | | **0.04** | 0.02 | | | |
| | | NM_198591.4 | 1485 | GAACT | **NT** | NT | | | **0.06** | 0.02 | | | |
| TPT1 | 45337310 | NM_001286273.2 | 874 | GGACT | **NT** | NT | **0.42** | 0.11 | **0.11** | 0.02 | **0.15** | **-** | **-** |
| | | NM_003295.4 | 709 | GGACT | **NT** | NT | | | **0.10** | 0.02 | | | |
| | 45337303 | NM_001286273.2 | 881 | AGACA | **NT** | NT | **0.12** | 0.07 | **0.08** | 0.02 | **0.04** | **-** | **-** |
| | | NM_003295.4 | 716 | AGACA | **NT** | NT | | | **0.04** | 0.02 | | | |
| | 45337294 | NM_001286273.2 | 890 | GGACT | **NT** | NT | **0.79** | 0.37 | **0.17** | 0.02 | **0.01** | **-** | **-** |
| | | NM_003295.4 | 725 | GGACT | **0.82** | 0.2 | | | **0.21** | 0.04 | | | |
| MRPL20 | 1402080 | NM_017971.4 | 529 | GGACT | **0.69** | 0 | **0.77** | 0.33 | **0.56** | 0.26 | **-** | **-** | **Y** |
| YTHDF2 | 28743593 | NM_016258.3 | 1504 | AGACT | **1** | 0.75 | **0.96** | 0.76 | **0.51** | 0.33 | **-** | **0.40** | **-** |
| ACTG1 | 81511529 | NM_001614.5 | 533 | GGACT | **1** | 0.23 | **0.92** | 0.34 | **0.41** | 0.12 | **-** | **0.84** | **-** |
| | | NM_001199954.3 | 652 | GGACT | **0.95** | 0.19 | | | **0.37** | 0.19 | | | |
| SEC11A | 84669674 | NM_014300.4 | 949 | AGACT | **NT** | NT | **0.22** | NT | **0.18** | 0.03 | **-** | **-** | **Y** |
| PARP1 | 226361173 | NM_001618.4 | 3496 | AGACT | **NT** | NT | **0.44** | 0.47 | **0.07** | 0.05 | **-** | **0.02** | **-** |

**Table S4 DNA probes used in the SELECT assay.**

| Name | Sequence |
|---|---|
| qPCR-A-F | 5'agatcgagagtgagtcgtgtgaat |
| qPCR-m6A-F | 5'agatcggaagagcgtagtgtatga |
| EMB1467_2533-Dp-A | 5'caccgcaagctaaacccgagatacaattcacacgactcactctcgatct |
| EMB1467_2533-Up-A | 5'cagcaggtgtgcaaattgcttataga |
| EMB1467_2629-Dp-m6A | 5'cctaataagaacaatacaagatgccatcatacactacgctcttccgatct |
| EMB1467_2629-Up-m6A | 5'gcttctacatgcaaatgtaaaaggg |
| PRP8A_7207-Dp-A | 5'cgatgctcctcgtgatagaactcctttattcacacgactcactctcgatct |
| PRP8A_7207-Up-A | 5'caggtgactccaggaaatgagtg |
| PRP8A_7340-Dp-m6A | 5'ctatactgcaaaataaagctaaatcactcatacactacgctcttccgatct |
| PRP8A_7340-Up-m6A | 5'gctcgccagtacaacatcttaca |
| CURT1B_969-Dp-A | 5'ccttcattccaattcatgaatggccattcacacgactcactctcgatct |
| CURT1B_969-Up-A | 5'cgcagctggtgttagattcttttga |
| CURT1B_1121-Dp-m6A | 5'cttgaggaatttacaacactttgactcatacactacgctcttccgatct |
| CURT1B_1121-Up-m6A | 5'gctgcgaccaaaacatctcatatat |
| NACA3_705-Dp-A | 5'cggatttaggtggtaagctccattcacacgactcactctcgatct |
| NACA3_705-Up-A | 5'agtagcagcatcaaagtaggaaaag |
| NACA3_802-Dp-m6A | 5'ccaaacgtatagtatataatgtcatacactacgctcttccgatct |
| NACA3_802-Up-m6A | 5'gcctcggcttctaaaatatgaaatg |
| RPL17B_589-Dp-A | 5'cttggacttggcagccaatatattcacacgactcactctcgatct |
| RPL17B_589-Up-A | 5'ttagagaaagaaagcttaagctgctga |
| RPL17B_688-Dp-m6A | 5'cttacaaaacgattcgagctaaagtcatacactacgctcttccgatct |
| RPL17B_688-Up-m6A | 5'cctcggcaagataaatctatccat |
